## Supplementary Material for "Structural basis for the activation and ligand recognition of the human oxytocin receptor"

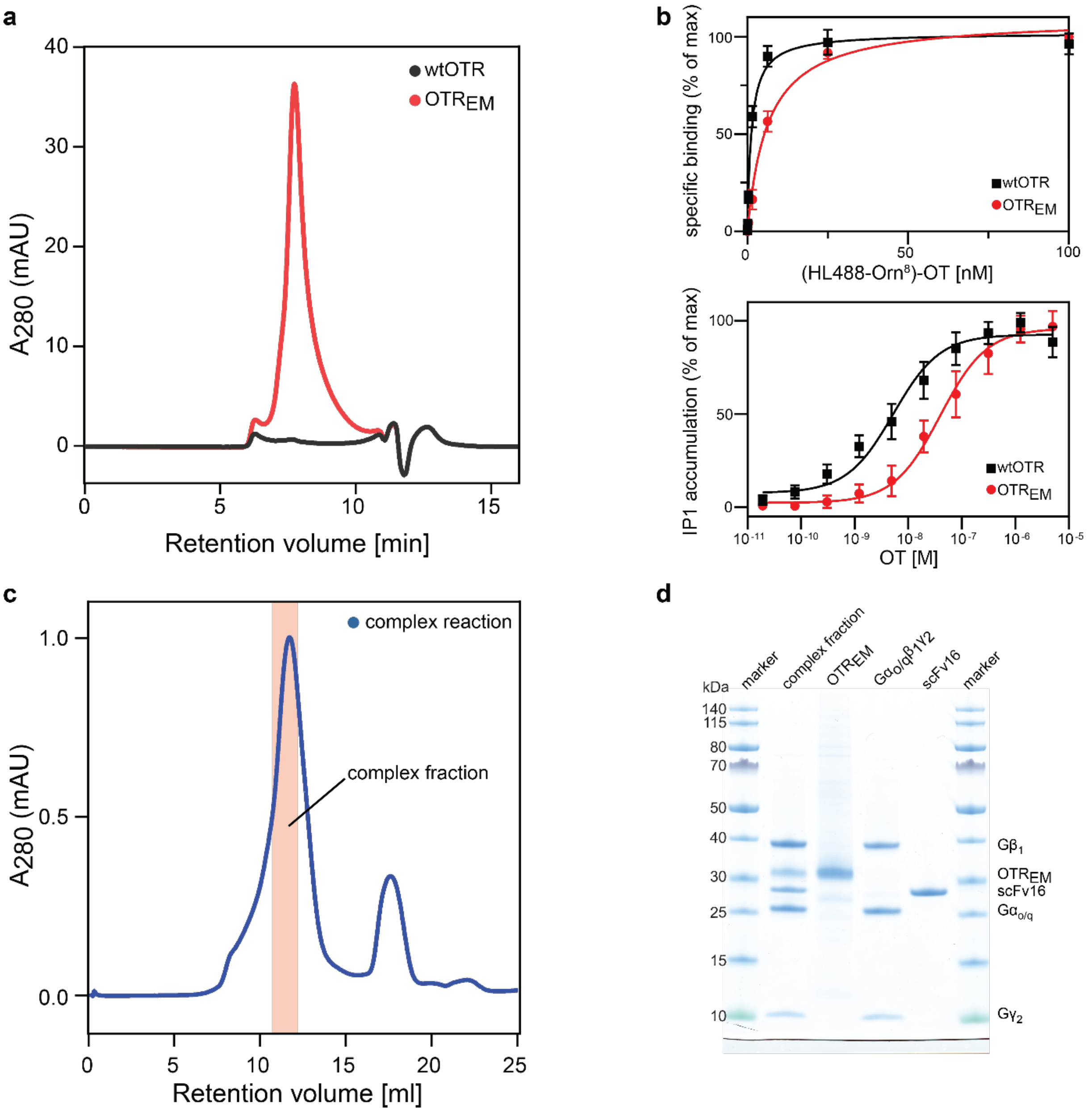


**Supplementary Fig. 1: Purification of $OTR_{EM}$ & complex formation**

**a** Small-scale analytical size-exclusion chromatography (SEC) profiles from initial purifications of wtOTR (black curve) and OTR-D153Y termed $OTR_{EM}$ (red curve). SEC profiles present fair loads. **b** Agonist profiles of wtOTR and $OTR_{EM}$. Dose-response curves were obtained from IP1 accumulation assays, and saturation binding assays were measured by whole-cell ligand binding assays. Curves are shown with standard deviation from at least two independent experiments performed in duplicates (IP1) and triplicates (ligand binding), respectively. **c** SEC profile of the OTR:OT:$G_{o/q}$:scFv16 complex. The red rectangle highlights the pooled fraction used for cryo-EM analysis. **d** LDS-PAGE gel of pooled complex fraction and the single components.

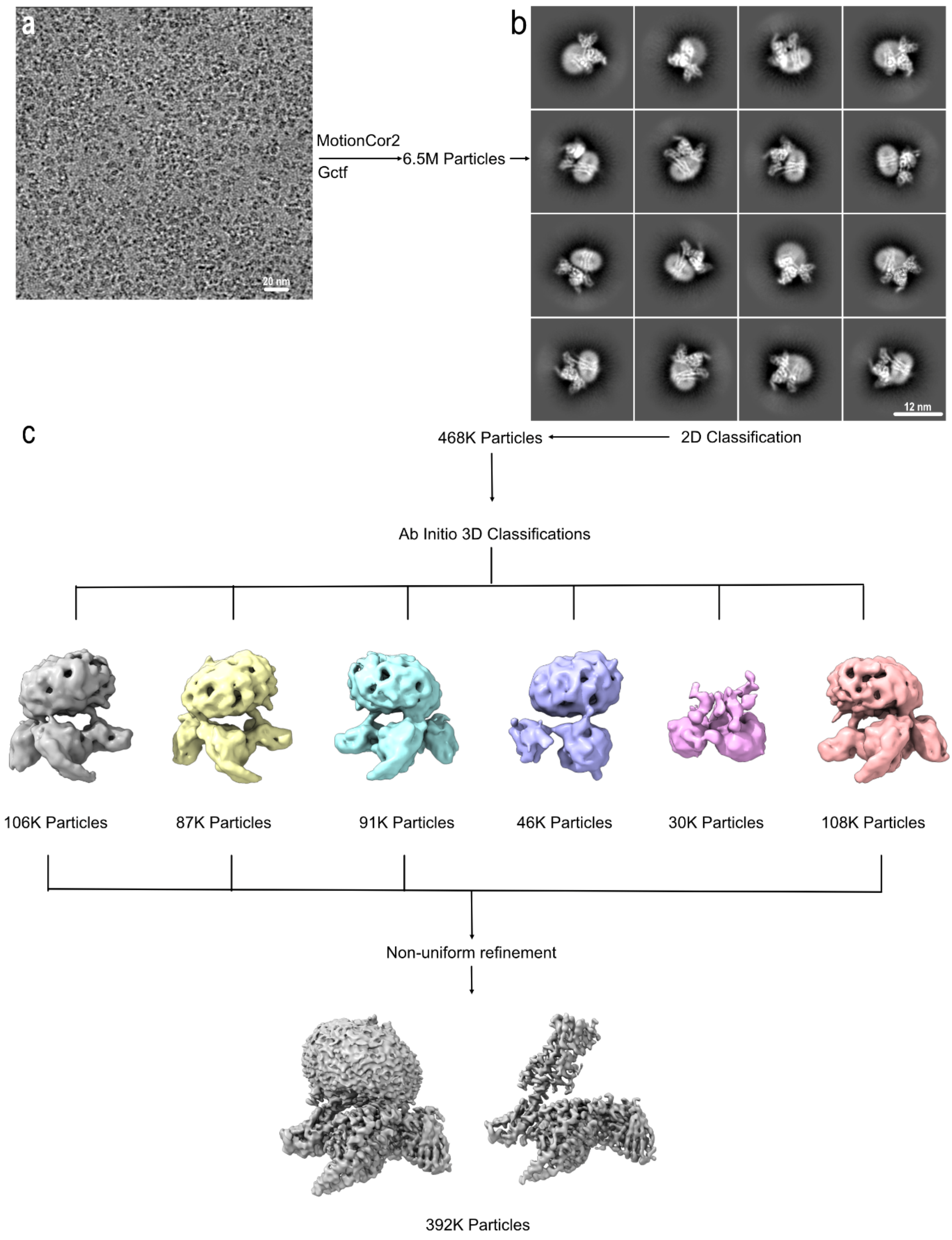


**Supplementary Fig. 2: Overview of single-particle cryo-EM data processing**

**a** Representative cryo-EM micrograph of the OTR:OT:$G_{o/q}$:scFv16 complex. Scale bar, 20 nm.

**b** Representative 2D averages showing distinct secondary structure features from different views of the complex. **c** 3D classification workflow and refinement.

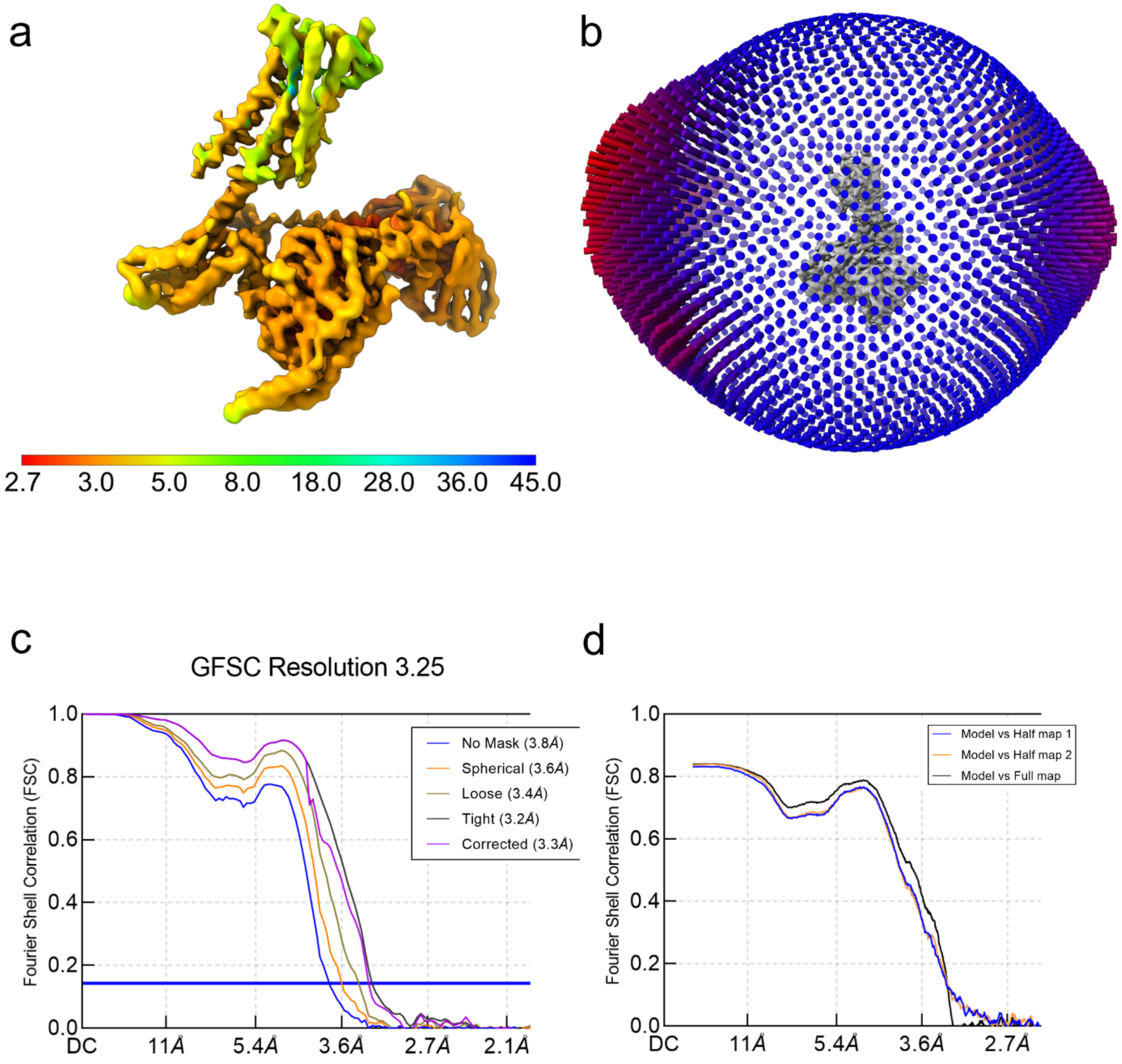


**Supplementary Fig. 3: Resolution of the OTR:OT:$G_{o/q}$:scFv16 complex.**

**a** Local resolution analysis of the OTR:OT:$G_{o/q}$:scFv16 complex. **b** Angular distribution of the particle orientations of the OTR:OT:$G_{o/q}$:scFv16 complex. **c** The gold-standard Fourier shell correlation curves for the map of the OTR:OT:$G_{o/q}$:scFv16 complex. **d** For cross-validation, FSC curves of the refined model versus full map (black), refined map versus half map 1 (blue), and refined model versus half map 2 (orange) were calculated.

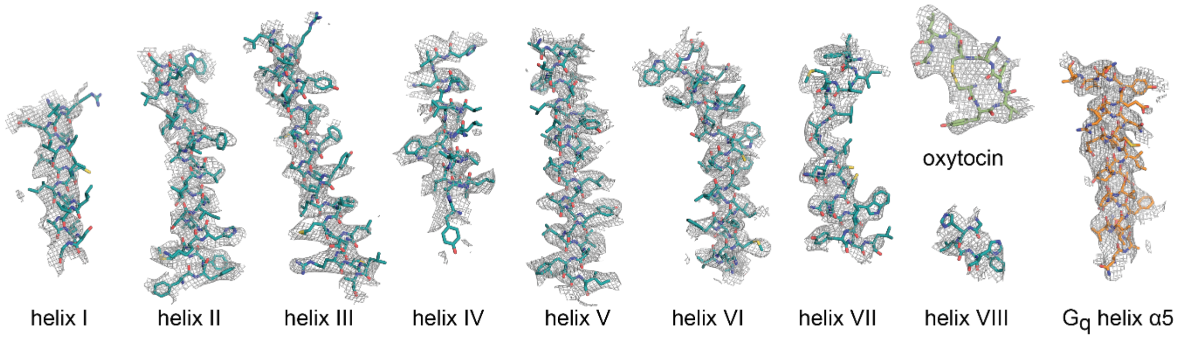


**Supplementary Fig. 4: Cryo-EM density within OTR.**

Cryo-EM density maps for all OTR transmembrane helices, helix VIII, oxytocin, and the interacting $G_q$ α5 helix of the G protein.

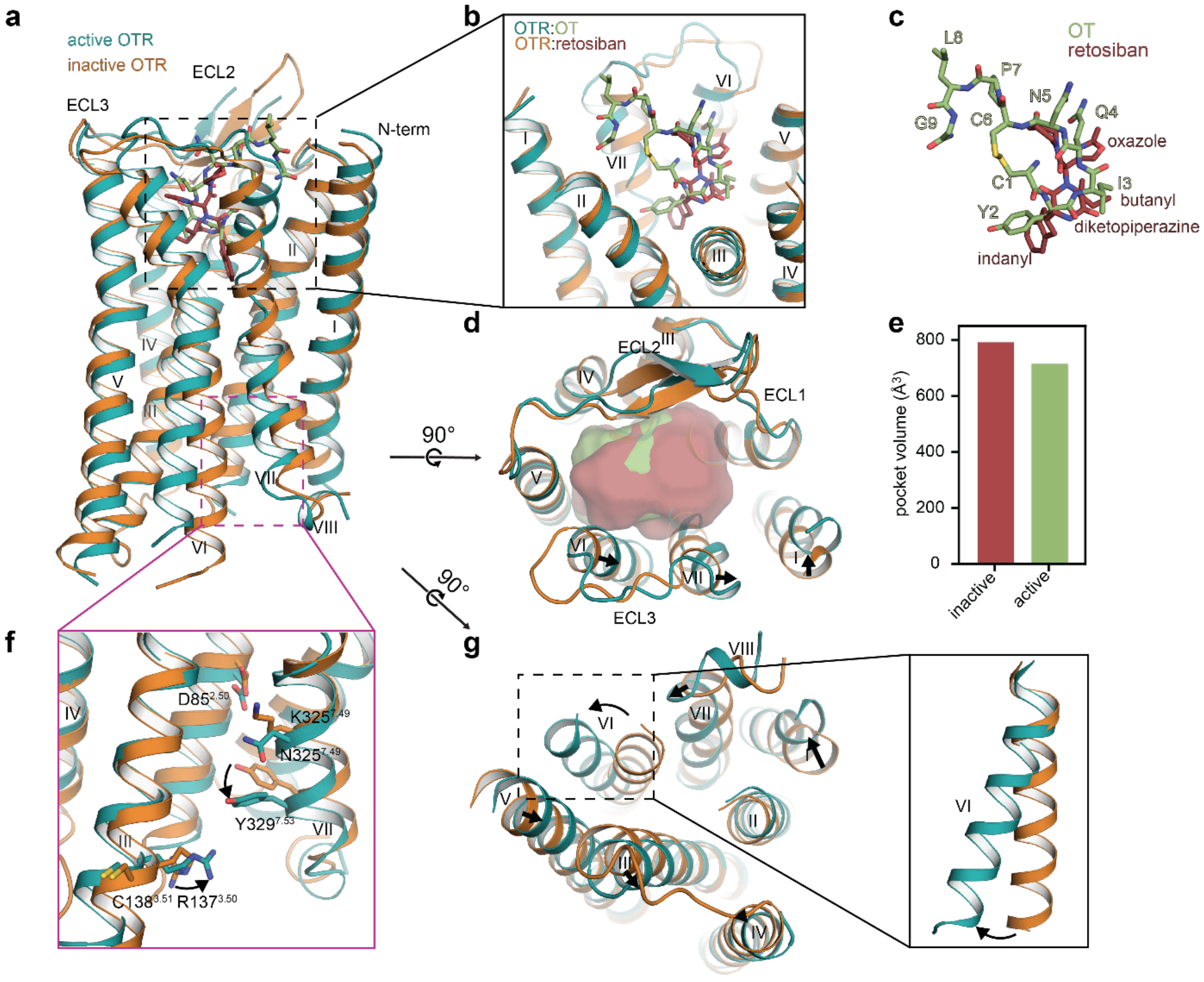


**Supplementary Fig. 5: Activation mechanism of the OTR.**

**a** Superposition of active OTR:OT complex (teal) and inactive OTR:retosiban complex (orange, PDB ID: 6TPK). **b** Close-up on binding pockets of OT and retosiban viewed from the extracellular side. **c** OT and retosiban binding modes as viewed from the membrane plane. **d** Extracellular view of the super-positioned receptors with calculated pocket volume shown as surface representation. Arrows indicate shifts of the extracellular helix tips from inactive to active state. **e** Calculated pocket volumes for inactive and active OTR conformations. Pocket volumes were calculated with POVME 2.0[1]. **f** Close-up view on class A-specific microswitch motifs DRC and NPxxY. Arrows indicate shifts of microswitch residues from inactive to active state. **g** Intracellular view of super-positioned receptors with additional close-up view on helix VI. Arrows indicate shifts of the intracellular helix tips from inactive to active state.

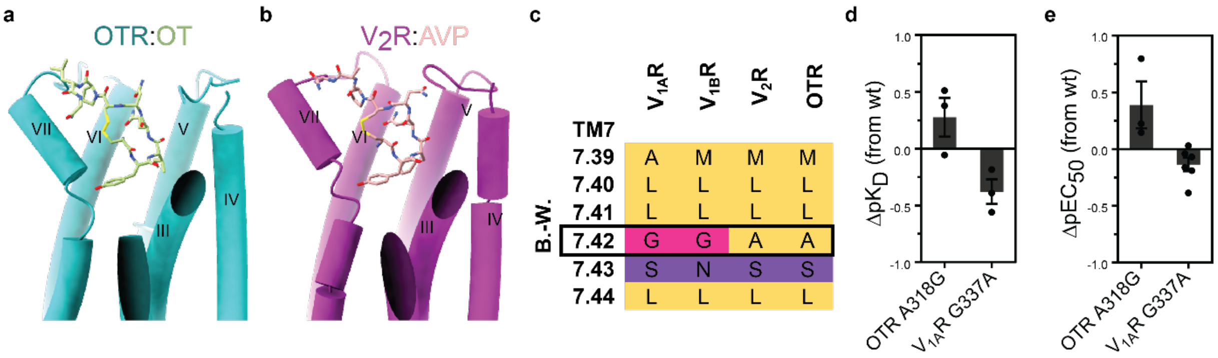


**Supplementary Fig. 6: Conserved activation mechanism by oxytocin and vasopressin.**

**a** Cylindrical representation of active OTR:OT complex with close-up on kink in helix VII. **b** Cylindrical representation of active $V_2R$:AVP complex (PDB ID: 7DW9) with close-up on kink in helix VII. **c** Amino acid sequence alignment of the kink region for all human oxytocin and vasopressin receptors. Amino acid positions are denoted in Ballesteros-Weinstein numbering (B.-W.)[2]. **d** OT affinity profiles of OTR and $V_{1A}R$ kink region mutants. Bars represent differences in affinity of the congnate ligand (mean $pK_D$ ± SEM from three independent experiments in triplicates) compared to wtOTR or wt$V_{1A}R$. **e** OT IP1 accumulation dose-response curves of OTR and $V_{1A}R$ kink region mutants. Bars represent differences in IP1 accumulation potency of the cognate ligand (mean $pEC_{50}$ ± standard deviation from at least three independent experiments in duplicates) compared to wtOTR or wt$V_{1A}R$.

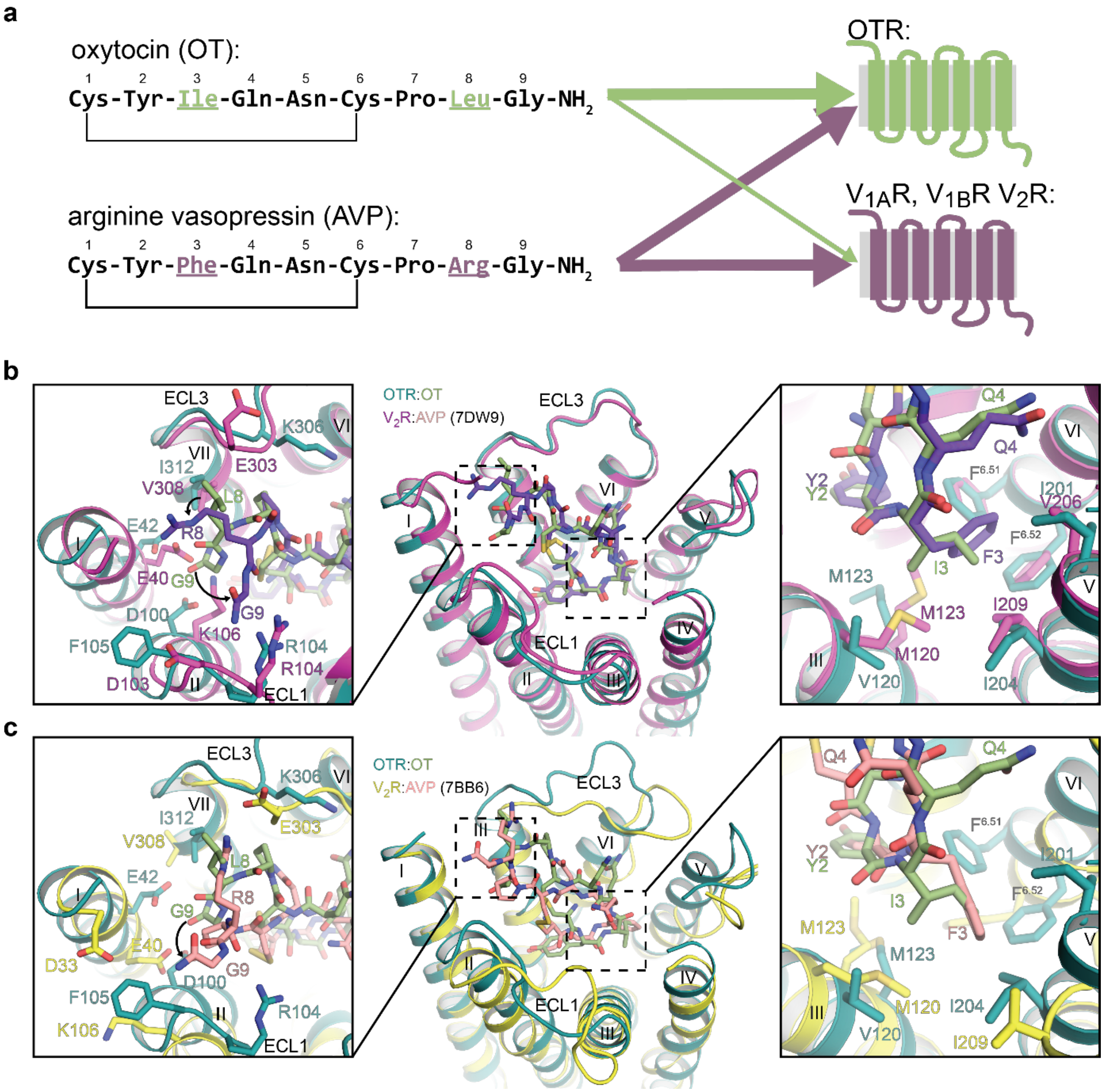


**Supplementary Fig. 7: Comparison of the OTR and V$_2$R orthosteric binding pockets.**

**a**. (left) Amino acid sequences of OT and AVP. Amino acid differences between the closely related hormones are highlighted. (right) Simplified specificity profile of OT and AVP for oxytocin and vasopressin receptors. Line thickness indicates affinity towards indicated receptors. **b,c** Structural superposition of OTR:OT with V$_2$R:AVP structures (b, PDB ID: 7DW9; c, PDB ID: 7BB6), illustrating the significant differences between AVP positions 3 and 8 in the two V$_2$R:AVP structures. Arrows indicate conformational changes in non-conserved positions of AVP and OTR. (left) Close-up on sub-pocket binding AVP/OTR position 8.

(middle) Overview of binding pockets. (right) Close-up on sub-pocket binding AVP/OTR position 3.

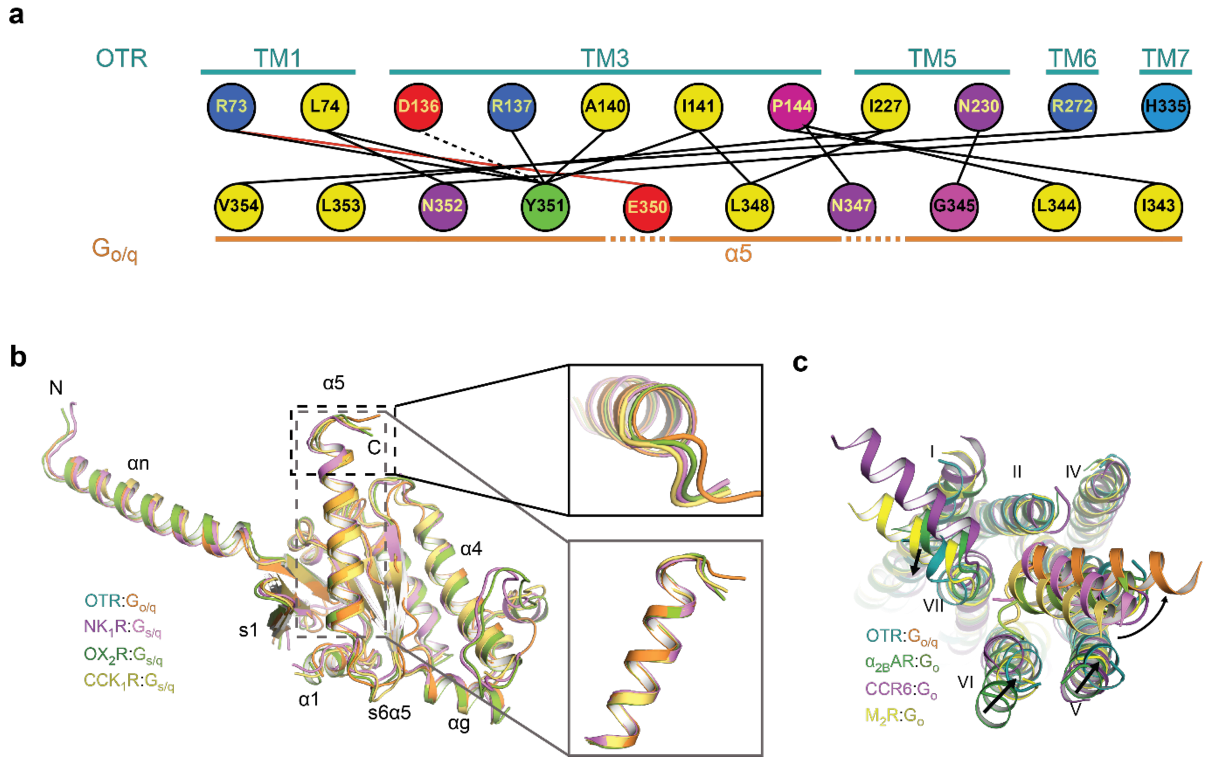


**Supplementary Fig. 8: Interactions of OTR and G$\alpha_{o/q}$**

**a** Schematic drawing of direct interactions between OTR and the α5 helix of G$\alpha_{o/q}$. Hydrogen bonds are drawn as dashed lines. Salt bridges are indicated by lines coloured in red. Amino acids are coloured according to their biophysical properties. **b** Structural superposition of OTR-bound $G_{o/q}$ with $G_{s/q}$ from different signalling complexes: $NK_1R$:$G_{s/q}$ (purple; PDB ID: 7P00), $OX_2R$:$G_{s/q}$ (green; PDB ID: 7L1U), and $CCK_1R$:$G_{s/q}$ (yellow, PDB ID: 7MBY) with two different close-up views on Gα5. **c** Structural superposition of OTR with $\alpha_1AR$:$G_o$ (green; PDB ID: 6K41), CCR6:$G_o$ (purple; PDB ID: 6WWZ), and $M_2R$:$G_o$ (yellow; PDB ID: 6OIK) as viewed from the intracellular side with a focus of receptor – G protein interaction. Arrows indicate structural differences between intracellular receptor helix tips and α5 orientations.

**Supplementary Table 1. Single-particle cryo-EM statistics**

| | OTR:OT:$G_{o/q}$:scFv16<br>PDB ID: 7QVM |
|---|---|
| **Data collection** | |
| Microscope | Titan Krios G3i |
| Detector | Gatan K3 |
| Energy filter slit width (eV) | 20 |
| Magnification | 130,000 |
| Voltage (kV) | 300 |
| Electron exposure ($e^-/Å^2$) | 63.7 |
| Defocus range (µm) | 0.8-2.4 |
| Pixel size (Å) | 0.65 |
| Symmetry imposed | C1 |
| Number of Micrographs | 10,276 |
| Initial particle images (no.) | 6.5 Mio |
| Final particle images (no.) | 392,369 |
| Map resolution (Å) | 3.25 |
| FSC threshold | 0.143 |
| **Refinement** | |
| Number of atoms | |
| All | 8,551 |
| Protein | 8,482 |
| Ligand | 69 |
| Model validation | |
| CC map vs. model (%) | 76 |
| RMSD | |
| Bond lengths (Å) | 0.27 |
| Bond angles (°) | 0.640 |
| Ramachandran statistics | |
| Favored regions (%) | 96.4 |
| Allowed regions (%) | 3.5 |
| Outliers (%) | 0.0 |
| Rotamer outliers (%) | 0.0 |
| C-beta deviations (%) | 0.0 |
| Clashscore | 11.4 |
| *MolProbity* overall score | 1.8 |

**Supplementary Table 2. Effects of mutations on OT-induced IP1-accumulation**

| construct | $EC_{50}$ [nM] | $\Delta pEC_{50}$ | $E_{max}$ (% of wt) | n |
|---|---|---|---|---|
| wtOTR | 8.1 ± 4 | - | 100 | 6 |
| $OTR_{EM}$ | 42.8 ± 17.1 | -0.64 ± 0.3 | 229 ± 29 | 2 |
| Q92A | 222.3 ± 141 | -1.72 ± 0.28 | 12 ± 1 | 3 |
| Q96A | 1540 ± 286.3 | -2.75 ± 0.05 | 68 ± 13 | 3 |
| K116A | 6.1 ± 0.8 | -0.36 ± 0.03 | 48 ± 5 | 3 |
| Q119A | 444.3 ± 192.6 | -2.15 ± 0.16 | 107 ± 33 | 3 |
| M123A | n.a. | - | (1 ± 1) | 3 |
| Q171A | 992.5 ± 180 | -2.56 ± 0.04 | 90 ± 12 | 3 |
| Q171N | 39.1 ± 10.2 | -1.14 ± 0.1 | 74 ± 11 | 3 |
| F175A | 3482 ± 857.7 | -3.09 ± 0.07 | 66 ± 12 | 3 |
| W188A | 157.2 ± 60.5 | -1.21 ± 0.31 | 86 ± 10 | 2 |
| I201A | 89.5 ± 15.4 | -1.52 ± 0.12 | 22 ± 7 | 3 |
| I204A | n.a. | - | (-2 ± 4) | 3 |
| F291A | 585.7 ± 223.3 | -1.78 ± 0.66 | 28 ± 7 | 2 |
| F292A | 4.6 ± 1.7 | 0.33 ± 0.66 | 9 ± 2 | 2 |
| Q295A | 52 ± 2.3 | -0.76 ± 0.46 | 32 ± 1 | 2 |
| L316A | 22.2 ± 10.4 | -0.34 ± 0.26 | 18 ± 4 | 2 |
| A318G | 4.2 ± 1.1 | 0.39 ± 0.21 | 50 ± 2 | 3 |
| $wtV_{1A}R$ | 158 ± 27.6 | - | 100 | 6 |
| G337A | 250 ± 90.4 | -0.14 ± 0.06 | 215 ± 15 | 6 |

HTRF-based measurements of IP1 accumulation in HEK293T cells expressing wild-type and mutated receptor variants. Activation curves were analyzed by fitting each experiment separately to a three-parameter logistic equation. All values are expressed as mean ± SEM of the indicated number of independent experiments performed in duplicate. n.a., no activation.

**Supplementary Table 3. Effects of mutations on OT binding**

| construct | $K_D$ [nM] | $\Delta pK_D$ | $B_{max}$ (% of wt) | n |
|---|---|---|---|---|
| wtOTR | 1.4 ± 0.2 | 0 | 100 | 6 |
| $OTR_{EM}$ | 6.4 ± 0.7 | -0.6 ± 0.1 | 252 ± 10 | 3 |
| A318G | 0.9 ± 0.2 | 0.28 ± 0.17 | 28 ± 5 | 3 |
| $wtV_{1A}R$ | 9.3 ± 2.2 | 0 | 100 | 3 |
| G337A | 20.9 ± 0.2 | -0.38 ± 0.11 | 165 ± 23 | 3 |

Whole-cell specific saturation binding experiment of fluorescently labelled peptide OT-HL488 to HEK293T cells expressing wild-type and mutated receptor variants. Binding curves were analyzed by fitting each experiment separately to a one-site saturation binding equation. All values are expressed as mean ± SEM of the indicated number of independent experiments performed in triplicate. $B_{max}$ values indicate the amount of functional receptor.
